# Structural adaptability and surface activity of tardigrade-inspired peptides

**DOI:** 10.1101/2023.10.27.564388

**Authors:** Giulia Giubertoni, Sarah Chagri, Pablo G. Argudo, Federico Caporaletti, Alessandro Greco, Leon Prädel, Alberto Pavan, Ioana M.Ilie, Yong Ren, David Ng, Mischa Bonn, Tanja Weil, Sander Woutersen

## Abstract

Tardigrades are unique micro-animals that withstand harsh conditions, such as extreme temperatures and desiccation. Recently, it was found that specific cytoprotective proteins are essential for ensuring this high environmental tolerance. In particular, cytoplasmic abundant heat soluble (CAHS) proteins, which are intrinsically disordered, adopt more ordered conformations upon desiccation, and are involved in the vitrification of the cytoplasm. The design and synthesis short peptides capable of mimicking the structural behavior (and thus the cytoprotective properties) of CAHS proteins would be beneficial for potential biomedical applications, including the development of novel heat-resistant preservatives for sensitive drug formulations. As a first step in this direction, we selected several model peptides of varying lengths derived from the conserved CAHS motifs 1 and 2, which are part of the intrinsically disordered CAHS-c2 region. We then studied their structures using circular dichroism and linear and two-dimensional infrared spectroscopy in the presence of the desolvating agent TFE (2,2,2-trifluoroethanol), which mimics desiccation. We found that the CAHS model peptides are mostly disordered at 0% TFE (a result that we confirmed by molecular dynamics simulations), but adopt a more α-helical structure upon the addition of the desolvating agent, similar to what is observed for full CAHS proteins. Additionally, we employed sum frequency generation to investigate the surface activity of the peptides at the air/water interface to mimic a partial dehydration effect. Interestingly, all model peptides are surface active and also adopt a helical structure at the air/water interface. Thus, the selected sequences represent promising model peptides that show similarities in the physicochemical behavior to full CAHS proteins. Our results also suggest that arginine might be a crucial element in defining the strong propensity of these peptides to adopt a helical structure. In the future, the use CAHS model peptides to design new synthetic peptide-based materials could make it possible to mimic and exploit the cytoprotective properties of naturally occurring tardigrade proteins.

**SIGNIFICANCE:** Tardigrades are micro-animals that can survive extreme conditions such as desiccation and high temperatures. Recent work has shown that this capability is related to the presence of specific proteins that can remodel in order to protect the organism’s cells. Mimicking this behavior using small peptides that preserve the structural properties of the full proteins is highly desirable in potential biomedical applications, such as the storage of heat-sensitive drugs. Here, we study the structural properties of model peptides derived from the conserved region of cytoplastic tardigrade proteins, and show that these peptides preserve some of the conformational behavior of the full protein under drying conditions. These peptides can therefore be used as a starting point for the design of synthetic model systems based on tardigrade-inspired peptides for tailored applications.

## INTRODUCTION

Tardigrades, also known as “water bears”, are microscopic organisms that can withstand harsh environmental conditions, such as high temperatures and desiccation.(1, 2) Research into the biomolecular mechanisms behind their high-stress tolerance has shown that certain proteins play an essential role in protecting the integrity of cellular components under extreme conditions.(3)

Three novel protein families have been discovered in the past decade that are conserved among different species of tardigrades: Cytoplasmic, Secreted, and Mitochondrial Abundant Heat Soluble (CAHS, SAHS, and MAHS) proteins.(3, 4) Although the cytoprotective mechanism of CAHS proteins is not fully understood, recent work from Boothby et al. has shown convincing evidence that CAHS proteins under desiccation undergo a liquid-to-gel transition, which restricts the molecular motions of both the protein itself and the desiccation-sensitive biological materials embedded within the gel, thereby promoting the maintenance of cell volume and the induction of reversible biostasis. Such gelation is connected to a structural change of the protein during the desiccation: CAHS proteins are intrinsically disordered proteins. Yet, upon desiccation, the central domain folds into an α-helix, whereas the external domains adopt a β-sheet structure. Helix-helix interactions between the proteins are crucial to drive the liquid-to-gel transition.(5)

Mimicking the structural adaptability of CAHS, SAHS and MAHS proteins to achieve comparable biological functionality has been a long-standing goal in the development of synthetic bionanomaterials. For the family of LEA (Late Embryogenesis Abundant) proteins, there have already been attempts to mimic their protective effect in preventing heat induced aggregation by using model peptides inspired by conserved LEA motifs. A recent example is the model LEA peptide (AKDGTKEKAGE)2, which can effectively inhibit heat-induced irreversible denaturation and inactivation of lysozyme in buffer solution.(6) Analogous to the research into LEA-inspired peptides, imitating some of the biological functions of CAHS proteins by using model peptides would be desirable for many applications, such as cryopreservation or heat-preservation of sensitive drug formulations. Therefore, we selected four peptide sequences inspired by conserved regions of CAHS proteins (Table 1) and studied their structural adaptability under desiccating conditions. We also investigated the effect of sequence length by synthesizing CAHS11aa as a truncated version of the longer model peptide CAHS22aa. As a negative control, we chose a peptide sequence inspired by a non-CAHS tardigrade protein, the secreted abundant heat-soluble protein 1 (SAHS1). Since the model peptide SAHS1C3 (which we will refer to as the control peptide) was not derived from the CAHS protein family, we hypothesized that its structural adaptability and surface activity would greatly differ from the CAHS model peptides.

**Table 1:** CAHS model peptides and SAHS-inspired control peptide: names, sequence, number of amino acids and protein of origin (PDB). Bold letters highlight the amino acids that are more favorable to participate in the formation of α-helix.(7)

| Name | Sequence | Number | PDB |
| --- | --- | --- | --- |
| CAHS1M1 | Y**R**QTEAE**A****E****K****I****R****R****E****L****E****K** Q | 19 | J7M799 (CAHS motif 1) |
| CAHS1M2 | **K** Q**K**K MIDVES**R****Y****A****K****K** DMD**R****E** | 20 | P0CU51 (CAHS motif 2) |
| CAHS11aa | **A****E** **K** **I****R****K****E****L****E****K****Q** | 11 | P0CU45 |
| CAHS22aa | **A****E** **K** **I****R****K****E****L****E****K** QHARDVE**F****R****K****S****L** | 22 | P0CU45 |
| SAHS1C3 (control) | **K** YTEDG**D****K****L****V****A** | 11 | P0CU40* (except E114→ D) |

To study the secondary structure of the model peptides in bulk and their changes upon mimicking drying conditions, we combine circular dichroism (CD) with linear and two-dimensional infrared spectroscopy (IR and 2D-IR, respectively). CD spectroscopy is a powerful tool to distinguish between random coil, α-helical and β-sheet structures as these have quite distinct spectral signatures.(8) IR spectroscopy also reports on protein conformation, providing additional and complementary structural information. The molecular vibrations of the amide groups, in particular the amide I mode (which mainly involves the carbonyl stretching) are sensitive to the protein conformation (9); in theory, the absorption frequency of the amide I mode can be used as a structural marker. However, the infrared absorption spectra in the amide I region are generally rather congested (which is also a problem in CD spectroscopy), limiting our ability to disentangle the underlying amide I bands in an unambiguous manner. This problem can be partially solved by using 2D-IR spectroscopy (10), where the absorption bands become narrower and, thus, allow a better disentanglement of the spectra. Moreover, the secondary structure of the model peptides can be characterized at the air/water interface, known for its high hydrophobicity,(11) which allows the emulation of a partial dehydration effect.(12) Through the application of sum frequency generation (SFG), we can discern molecular order due to symmetry selection rules(13), and detect distinct characteristic frequencies associated with various molecular vibrational modes as it offers chemical specificity(14). This enables us to determine the amide I region structure of the first two layers of peptides, and thus their secondary structure.

We find that all model peptides inspired by the conserved domain of the CAHS proteins show a similar structural change upon drying compared to the full protein: under hydrating conditions the peptides have an unordered structure, whereas upon dehydrating they adopt a helical conformation. We note that the control peptide, expected to have different physicochemical properties, has a more ordered structure, consisting of a bend/turn that is stabilized under drying conditions. Furthermore, sequence-structure analysis suggests that specific patterns in the sequence, such as alternating of positive and negative charges, might be crucial in stabilizing alpha-helical conformations, as suggested in literature(15, 16). Finally, our observations indicate that the four model peptides not only exhibit surface activity but also, consistent with bulk findings, adopt a helical structure under partially dehydrated conditions, suggesting that this conformation is the most preferred in these circumstances.

## MATERIALS AND METHODS

### Peptide synthesis

CAHS11aa was synthesized using the Fmoc solid phase peptide synthesis strategy by Merrifield, synthesizing the peptide from C to N-terminus in a microwave-assisted peptide synthesizer. Fmoc–Gln (Trt) preloaded Wang resin (100–200 mesh) (0.1 mmol) was swollen in DMF for 1 h before use. First, the Fmoc group was cleaved by two deprotection steps using 20% v/v piperidine in DMF (3 mL) for 2 and 5 min at 75 °C. Fmoc-Lys(Boc), Fmoc-Glu(OtBu), Fmoc-Leu, Fmoc-Arg(Pbf), Fmoc-Ile and Fmoc-Ala (5 equiv in 2.5 mL) were coupled for 20 min using DIC (5 equiv in 1 mL) and Oxyma (10 equiv) in 0.5 mL DMF The peptide was purified by HPLC using the Atlantis T3 column at a 10 mL/min flow rate. The gradient started at a 95:5 ratio of water:ACN (+0.1% TFA) and was kept constant for 1 min, after which the ACN content was increased to 30% in 20 min. The retention time of CAHS11aa was 3.0 min. The peptide was received in a yield of 7.1% (21 mg). CAHS22aa was synthesized using the Fmoc solid phase peptide synthesis strategy by Merrifield, synthesizing the peptide from C to N-terminus in a microwave-assisted peptide synthesizer. Fmoc–Leu preloaded Wang resin (100–200 mesh) (0.1 mmol) was swollen in DMF for 1 h before use. First, the Fmoc group was cleaved by two deprotection steps using 20% v/v piperidine in DMF (3 mL) for 2 and 5 min at 75 °C. Fmoc-Lys(Boc), Fmoc-Glu(OtBu), Fmoc-Leu, Fmoc-Arg(Pbf), Fmoc-Ile, Fmoc-Ala, Fmoc-His(Trt), Fmoc-Asp(OtBu), Fmoc-Val, Fmoc-Phe and Fmoc-Ser(tBu) (5 equiv in 2.5 mL) were coupled for 20 min using DIC (5 equiv in 1 mL) and Oxyma (10 equiv) in 0.5 mL DMF. The peptide was purified by HPLC using the Atlantis T3 column at a 10 mL/min flowrate. The gradient started at a 95:5 ratio of water:ACN (+0.1% TFA) and was kept constant for 1 min, after which the ACN content was increased to 30% in 20 min. The retention time of CAHS22aa was 1.5 min. The peptide was received in a yield of 2.0% (1.5 mg). CAHS1M1, CAHS1M2 and SAHS1C3 (control) were purchased via custom peptide synthesis from Jiangsu GenScript Biotech Co., Ltd. and received at ≥95% purity.

### Circular dichroism

CD spectra were recorded on a JASCO J-1500 spectrometer in a 0.1 cm High Precision Cell by HellmaAnalytics. The recorded data were processed in Spectra Analysis by JASCO and OriginPro9. For the preparation of the peptide solutions, each model peptide was dissolved in phosphate buffer (PB, 10 mM, pH 7.4) or a mixture of phosphate buffer and TFE (20, 50, and 70,% respectively) to achieve a peptide concentration of 0.1 mg/mL in each sample. The samples were subsequently measured and the spectra were recorded at wavelengths from 260 to 180 nm with a bandwidth of 1 nm, data pitch of 0.2 nm and scanning speed of 5 nm/min. Spectra were measured three times and accumulated.

### Infrared and two-dimensional infrared spectroscopy

A Perkin-Elmer Spectrum-Two FTIR spectrometer (resolution 2 cm^−1^) was used to measure the FTIR spectra of the background and peptide solutions. A detailed description of the setup used to measure the 2DIR spectra can be found in ref. 17. Briefly, pulses of wavelength 800 nm and with a 40 femtosecond duration are generated by using a Ti:sapphire oscillator, and further amplified by using a Ti:sapphire regenerative amplifier to obtain 800 nm pulses at 1 kHz repetition rate. These pulses are then converted in an optical parametric amplifier to obtain mid-IR pulses (∼ 20 μJ, ∼6100 nm) with a full width at half max (FWHM) of 150 cm^−1^. The beam is then split into a probe and reference beam (5% each), and a pump beam (90%) that is aligned through a Fabry-Pérot interferometer. The pump and probe beams are overlapped in the sample in an ∼250-μm focus. The transmitted spectra of the probe (*T*) and reference (*T* _0_) beams with the pump on and off are then recorded after dispersion by an Oriel MS260i spectrograph (Newport, Irvine, CA) onto a 32-pixel mercury cadmium telluride (MCT) array. The probe spectrum is normalized to the reference spectrum to compensate for pulse-to-pulse energy fluctuations. The 2DIR signal is obtained by subtracting the probe absorption in the presence and absence of the pump pulse. Parallel and perpendicular 2DIR spectra are recorded by rotating the pump beam at a 45^?^ angle with respect to the probe beam and selecting the probe beam component that is either perpendicular or parallel to the pump beam using a polarizer after the sample. To minimize pump-scattering contributions, we measured the average of two photoelastic-modulator-induced pump delays, such that the time delay between the scattered pump beam and the probe beam differs by 1/2 optical cycle in one delay with respect to the other. For all IR and 2D-IR measurements at 0% TFE, peptides were dissolved in phosphate buffer (10 mM, pH 7.4) at a concentration of 10 mg/ml, and the pH was adjusted to neutral by adding an adequate amount of NaOH. For all IR and 2D-IR measurements at 50 % of deuterated TFE (Sigma Aldrich), a peptide concentration of 5 mg/ml was used.

### Sum frequency generation spectroscopy

The SFG laser system consists of a seed and pump laser, a regenerative amplifier, and an optical parametric amplifier (OPA). Pulses with a 40 femtosecond duration are generated by using a Ti:sapphire seed laser (Mai Tai, Spectra Physics), stretched in time, and the selected ones are directed into a regenerative amplifier (Spitfire Ace, Spectra Physics). The Ti:Sapphire crystal in the amplifier is pumped with an Nd:YLF laser (Empower 45, Spectra Physics). After amplification, the selected pulses were guided through a compressor, which effectively restored them to their original 40 femtosecond duration. The output beam (1 kHz repetition rate, ∼800 nm wavelength) is split into two paths. The first path goes to an etalon (SLS Optics) to produce a spectrally narrow visible beam (FWHM ∼15 cm^−1^). The second path is used to pump an OPA (TOPAS-C, Light Conversion), in which the signal and idler beams are generated. The broadband infrared beam is generated by difference frequency generation from these beams. Finally, the SFG signal is detected by an electron-multiplied charge-coupled device camera (Newton EMCCD 971P-BV, Andor Technology). The polarization state of the SFG, Vis, and IR beams is controlled by polarizers and half-wave plates. In all experiments, SSP (s-polarized SFG, s-polarized VIS, p-polarized IR) polarization combination was used (where p/s denotes the polarization parallel/perpendicular to the plane of incidence defined by the direction of incidence of the beam and the perpendicular to the interface). Each experiment was performed using PBS buffer as the subphase under neutral pH conditions, a final peptide concentration of 0.09 mg/mL after injection, and a minimum time of 40 min. The samples were constantly rotated at ∼6 rpm, kept in a closed box, and subjected to a continuous flow of nitrogen to prevent absorption of infrared radiation by air. To process each SFG spectrum, the background-corrected sample spectrum was divided by the background-corrected reference spectrum to correct for the spectral shape of the IR beam. The background was recorded by blocking the IR beam. A z-cut quartz crystal was used as the reference. Each sample spectrum was recorded for 10 minutes, while the reference spectrum was for 10 seconds. The SFG spectra were fitted with Lorentzian peak shapes, following the procedure described elsewhere(13).

### Molecular dynamics simulations

Ten different initial conformations were generated for the CAHS11aa and the control SAHS1C3 peptide. Five peptides were generated using Alphafold2 (18), which predicted helical conformations for CAHS11aa with high confidence, and disordered structures for SAHS1C3. Five peptides per system were reconstructed using Monte Carlo sampling. More precisely, the residues were built initially in an excluded volume-obeying manner using CAMPARIv3 (http://campari.sourceforge.net) and then subjected to 1000000 elementary Monte Carlo steps (19, 20). This procedure randomizes dihedral angles hierarchically and guarantees that there is no spurious correlation between the starting models. All simulations were carried out using the GROMACS 2019.4 simulation package (21, 22) and the CHARMM36m force field (23). Four simulation systems were prepared consisting of a peptide (CAHS11aa or SAHS1C3) in 0M and 7M TFE. Ten independent simulations were carried out for each system for a total sampling time of 10 μs. The N- and C-termini of all fragments were capped with ACE and NME groups, respectively. For the systems with TFE, the molecules were added around the peptide prior to solvation. Each complex was then solvated in a cubic box (edge length of 6.9 nm) with TIP3P water molecules (24) to which 150 mM NaCl were added, including neutralizing counterions. Following the steepest descent minimization, the systems were equilibrated under constant pressure for 5 ns, with position restraints applied on the heavy atoms of the proteins. The temperature and pressure were maintained constant at 300 K and 1 atm, respectively, by using the modified Berendsen thermostat (0.1 ps coupling)(25) and barostat (2 ps coupling) (26). The production simulations were performed in the NVT ensemble in the absence of restraints. Given that the initial conformations of the peptides have been artificially generated, the first 500 ns of the NVT runs were considered as equilibration and only the subsequent 500 ns were used for the analysis. The short-range interactions were cut off beyond a distance of 1.2 nm, and the potential smoothly decays to zero using the Verlet cutoff scheme. Periodic boundary conditions were used and the Particle Mesh Ewald (PME) technique (27) was employed (cubic interpolation order, real space cutoff of 1.2 nm and grid spacing of 0.16 nm) to compute the long-range electrostatic interactions. Bond lengths were constrained using a fourth-order LINCS algorithm with 2 iterations (28). In all simulation,s the time-step was fixed to 2 fs and the snapshots were saved every 50 ps.

## RESULTS AND DISCUSSION

### Bulk structure

#### Hydrating condition

We first study the bulk structure of the tardigrade-inspired peptides in buffer solution by using circular dichroism (CD) and (2D-)IR spectroscopy. We combine these methods because CD is sensitive to helical structure(8), while IR spectroscopy can discriminate between other secondary structures(9, 29) and is sensitive to changes in the hydrating conditions. However, both IR and CD spectroscopy are strongly affected by spectral congestion, limiting our ability to disentangle the presence of specific spectral signatures that can uniquely be assigned to distinct secondary structures. Although different data analysis methods have been proposed to solve this issue, identifying secondary structures based only on the CD spectral shapes or IR absorption frequencies obtained via fitting may not be sufficient to determine the secondary structures present. For instance, an assignment based on the absorption frequency is not always unique because of secondary effects, such as solvent interactions, that may shift the amide I vibrational frequencies, and more specific spectral signatures are required. This problem can be partially overcome by 2D-IR spectroscopy.(30) The combination of CD and (2D-)IR spectroscopy is thus a powerful tool to characterize the structure of the peptides.

The CD spectra are shown in Fig. 1A. We observed that CAHS1M1 shows a positive signal at 190 nm, followed by two negative peaks around 210 and 225 nm. Such spectrum indicates an alpha-helical structure(8), and was observed for full CAHS proteins(31). Interestingly, the CD spectra of CAHS1M2, CAHS11aa, and CAHS22aa differ significantly from CAHS1M1.

**Figure 1:**
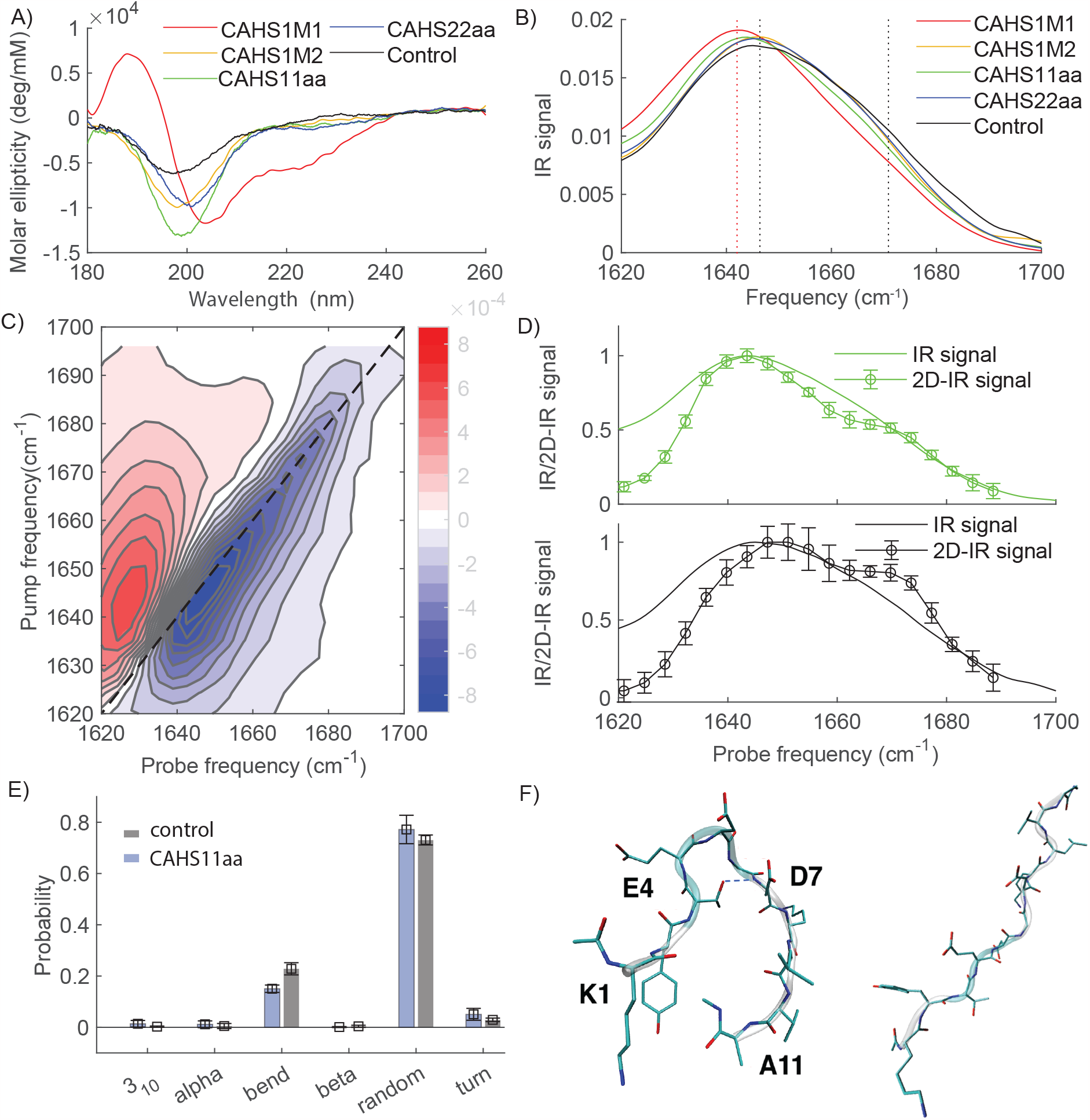
A) CD spectra of the model peptides at 0% TFE at a concentration of 0.1 mg/ml in in phosphate buffer (10 mM, pH 7.4). B) IR spectra of the model peptides at 0% TFE at a concentration of 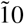 mg/ml in in phosphate buffer (10 mM, pH 7.4). Isotropic 2D-IR spectrum of CAHS11aa at a waiting time, Tw=1 ps. D) Comparison between the IR and 2D-IR diagonal slices of the bleach signals of the isotropic 2D-IR spectra of CAHS11aa (top) and the control peptide (bottom). E) Structural probability distributions for CAHS11aa and control peptide. F) Structural snapshots of the control peptide adopting a bend/turn structure (left), stabilized by a hydrogen bond between E and D amino-acids, and of the CAHS11aa (right).

These three peptides show a negative peak around 200 nm and a less intense one at 220 nm. This spectral signature can be mostly assigned to a random coil (unordered) structure.(8) Compared to CAHS11aa and CAHS1M2, CAHS22aa shows the minimum at a slightly higher wavelength and a slightly more pronounced negative peak at 220 nm, suggesting a greater propensity for alpha-helical structure. The control peptide shows a negative peak around 200 nm, which indicates that the SAHS peptide preferentially adopts an unordered structure.

Fig. 1B shows the IR spectra for the different peptides. The spectrum of CAHS1M1 is centered at around 1642 cm^-1^ with a shoulder around 1670 cm^-1^. We observe that the spectra of CAHS1M2, CAHS11aa, CAHS22aa and the control all show a similar spectral shape: a main band around 1648 cm^-1^, and again a shoulder at higher frequency around 1670 cm^-1^. In these spectra, it is challenging to disentangle the different vibrational bands that are present; and uniquely identify their vibrational frequencies, which can be used to characterize the peptide secondary structures.

To overcome these problems, we use two-dimensional infrared spectroscopy (2D-IR). Compared to the linear infrared spectra, 2D-IR better resolves the individual vibrational bands because the 2DIR signal scales quadratically with the absorption cross-section (while the conventional IR signal scales linearly with the cross-section), leading to narrower peaks and a removal of the solvent background(30, 32) In pump-probe 2D-IR spectroscopy, an intense, narrow-band infrared pump pulse (with adjustable center frequency ν_pump_) resonantly excites at a specific frequency (in the present case, in the amide I band). A delayed, broad band probe pulse probes the frequency-dependent IR-absorption change Δ*A*, which results from the excitation by the pump beam. Measuring the Δ*A* spectra for a range of ν_pump_ values, we obtain 2-dimensional spectra showing the pump-induced absorption change Δ*A*(ν_pump,_ ν_probe_) as a function of the pump and probe frequencies ν_pump_ and ν_probe_.(10) Fig. 1C shows the 2D-IR spectrum of CAHS11aa (2D-IR spectra of the other peptides are shown in the SI). When the pump frequency is resonant with the ν = 0 → 1 frequency of the amide I mode, part of the molecules are excited to the ν = 1 state of this mode, causing a decrease in the absorption at the ν = 0 → 1 frequency and an increase in absorption at the ν = 1 → 2 frequency (which is at a slightly lower value than the ν = 0 → 1 frequency due to the anharmonicity of the vibrational potential), resulting in a Δ*A* < 0 feature on the diagonal and a Δ*A* > 0 feature slightly to the left of the diagonal. In this way, each normal mode gives rise to a +/− doublet on the diagonal of the 2D spectrum (diagonal peaks), where the peaks colored in blue represent decreases in absorption (Δ*A* < 0) due to depletion of the amide-I ν = 0 state, and the signal at lower probe frequency colored in red represents the induced absorption of the ν = 1→2 transition. The diagonal peak is centered at 1640 cm^-1^, showing a shoulder at a higher frequency. Taking the diagonal slice of the 2D-IR spectrum (Fig. 1D), we observe that the diagonal slice now clearly shows two distinguished vibrational bands, one centered at 1645 and the other at 1670 cm^-1^. Similarly, we better resolve the vibrational bands underlying the IR spectrum of the control peptide (Fig. 1E). Interestingly, we find that the band at 1675 cm^-1^is more pronounced in the control peptide than in the CAHS11aa (which was not clearly visible in the IR spectra). A complete comparison between the diagonal slices of all model peptides is shown in the SI (Fig. S1). These two bands can be assigned to random coil (1648 cm^-1^) and bend/turn (1675 cm^-1^) structures(9). In the IR spectra, CAHS1M1 shows a main band at a lower frequency with respect to the other peptides, which we assign to an alpha-helical structure based on the CD results. These results indicate that three of our chosen model peptides favor a more unordered structure in bulk conditions with a small propensity to adopt a helical structure, and one (CAHS1M1) is already mostly helical in hydrating conditions. Although the control peptide also prefers random coil structure, the relatively higher intensity of the bend/turn peak indicates a stronger propensity to form a bend/turn structure than the other model peptides. The molecular dynamics simulations confirm that the bend/turn structures are more frequently populated by the control peptide (Fig. 1E). Specifically, the residue-based analysis shows that the central stretch ^5^DGD^7^ of the SAHS peptide attains bend/turn conformations over 50% of the simulation time (adopting the conformation shown in Fig. 1F), which is significantly less for the flanking residues in the peptide (Fig. S2). In contrast, the central stretch of the CAHS11aa peptide ^5^RKE^7^ attains bend/turn conformations ca. 20% of the simulation time, which is comparable with the neighboring residues. (Fig. 1F)

### Dehydrating condition

We then study conformational changes induced by water-deficient conditions by doing a titration with the desolvating agent TFE (2,2,2-trifluoroethanol) in order to mimic desiccation. First, we measure the CD spectra of peptides by increasing the TFE content from 0 to 20, 50, and 70%. These experiments showcase the specific properties of the CAHS model peptides, which are comparable in terms of folding as the above-mentioned LEA model peptides.(33) As the level of the desolvating agent TFE was increased from 20%, 50% to 70%, the CD signals at 190 nm and 208 nm increased significantly, indicating an increase in α-helical content of the peptide secondary structure of CAHS model peptides (Fig. 2). In contrast, the control peptide showed no significant changes in secondary structure, even at 70% TFE content (bottom in Fig. 2).

**Figure 2:**
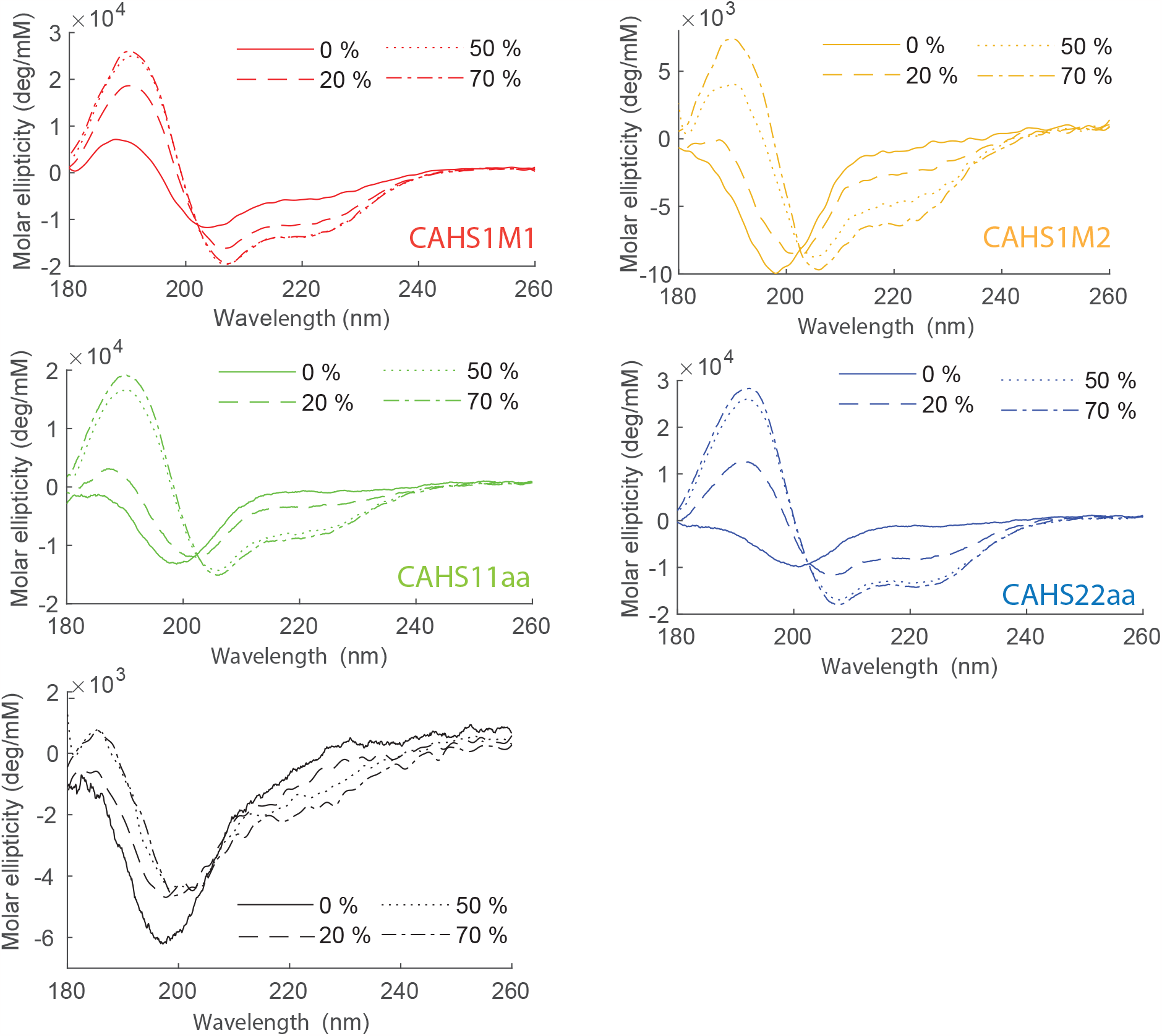
CD spectra of the model peptides dissolved at a concentration of 0.1 mg/ml in in phosphate buffer (10 mM, pH 7.4) while increasing the TFE concentration.

To obtain more structural detail, we perform 2D-IR at 50% of TFE content for CAHS11aa and control peptide (conventional IR spectra are shown in SI, Fig. S3). Fig. 3A shows the 2D-IR spectrum of CAHS11aa at 50% of TFE. Compared to the 2D-IR spectrum in Fig. 1C, the main peak is now shifted to higher frequency. In the diagonal-slice spectrum, the main band also shifts to higher frequency (from 1645 to 1655 cm^-1^) in the presence of TFE. A similar shift was observed in the case of LEA protein, where the absorption peak blueshifts from 1648 to 1660 cm^-1^ upon adding TFE because of the formation of helical structure.(34) Based on these previous results, we can assign the band at 1655 cm^-1^ to alpha-helical structure, indicating that upon increasing TFE content this peptide adopts a helical structure (as also corroborated by the CD data). Fig3B also shows a comparison between the diagonal slices of the control peptide, which did not show any significant change in the CD spectra when increasing the TFE concentration. The diagonal slices show that the band at 1670 cm^-1^increases in intensity, suggesting that drying increases the propensity of the control peptide to form a bend/turn structure.

**Figure 3:**
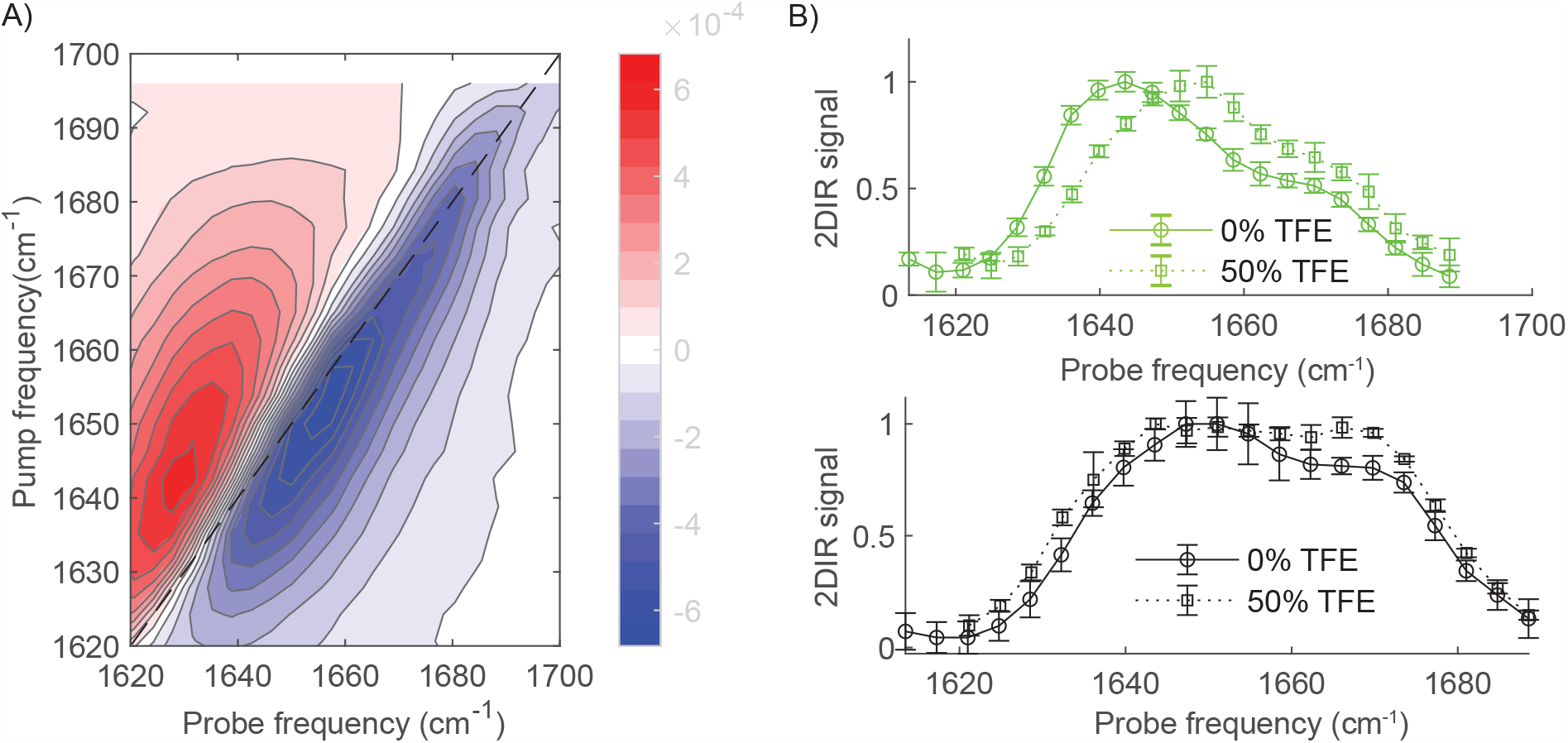
A) 2D-IR spectrum of CAHS11aa at 50% TFE concentration. B) Comparison of the 2D-IR diagonal slices of the bleach signals of the isotropic 2D-IR spectra of CAHS11aa and control peptides at of 0 and 50 % TFE concentration.

### Structure at the air/water interface

Finally, we characterize the behavior of the peptides at the air/water interface. Due to its hydrophobic nature, this interface also mimics desiccation.(11) In order to characterize the peptide structure, we conduct amide I SFG spectroscopy on the model peptides. SFG provides information about the structure and the orientation of interfacial proteins without interference from bulk molecules. Primarily arising from the carbonyl stretch of the amide bond, the amide I vibrational response is related to the peptide backbone.(29) Moreover, employing an SSP (s-polarized SFG, s-polarized VIS, p-polarized IR) polarization, the obtained spectra solely present the backbone amide I band, effectively eliminating side chain contributions and allowing an unobstructed analysis of the peptide’s reorientation.(35) It is important to note that the SFG signal intensity is determined not only by the number of proteins on the surface, but is also influenced by their orientation at the air/water interface. Specifically, the SSP geometry is sensitive to functional groups positioned perpendicular to the sample surface.(36) Fig. 4 shows the acquired SFG spectra, while the fitting parameters can be found in the SI. Since only ordered structures are visible in the SFG spectra, the presence of the amide I mode implies that the CAHS peptides occupy the interface and form a well-aligned layer at the air/water interface. Furthermore, consistent with prior CD and 2D-IR results under drying conditions, these peptides predominantly adopt an α-helical conformation. This is evidenced by a peak at ∼1645 cm^−1^, characteristic of helical structures.(35, 37–39) The mode at ∼1720 cm^−1^, typically attributed to the carbonyl vibrations of amino acid side chains,(40) can be assigned to protonated glutamic acid and aspartic acid side chains of the peptides.(41) Both amino acids can assume a protonated state within the peptide when present in a neutral pH environment.(42–44) In contrast, the control peptide does not display interfacial activity at the measured concentration, yielding a spectrum indistinguishable from buffer subphase. Finally, a broad low-intensity spectral feature is observed over the entire range of the amide I region corresponding to the H-O-H bending mode of the water.

**Figure 4:**
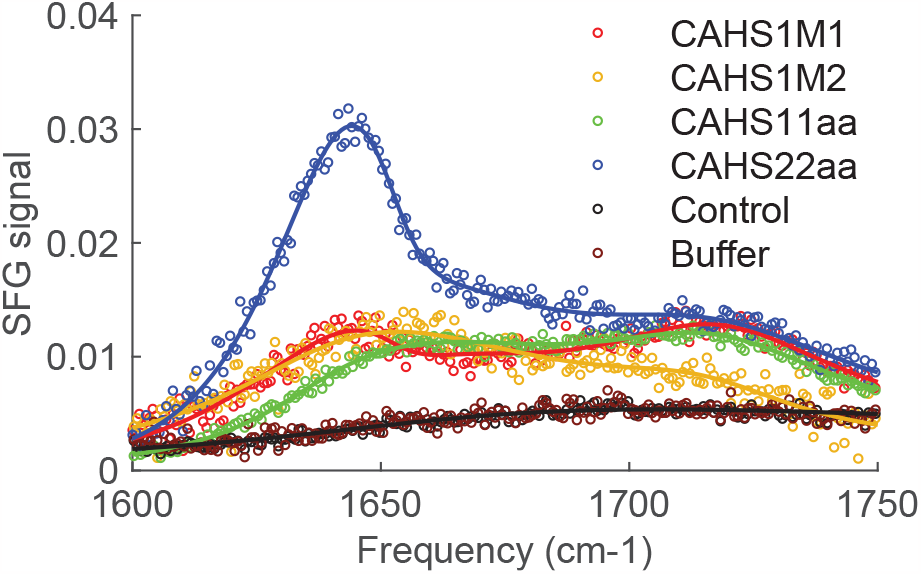
SSP amide I SFG spectra (open symbols) along with fits (solid lines) for the 5 model peptides and the buffer used at the air/water interface.

### Discussion and conclusion

We now compare the structure of the model peptides derived from the conserved region of CAHS protein and 1 control peptide in hydrating and dehydrating conditions. From the CD and (2D-)IR we find that the structure of the 3 model peptides (CAHS1M2, CAHS11aa and CAHS22aa) and the control one is mostly unordered, with different preferences for adopting a transient helical structure. Interestingly, CAHS1M1 adopts mainly a helical configuration already at 0 % TFE. Upon mimicking desiccation (either by adding TFE or by exposing the peptides to a water/air interface), we observed that all model peptides adopt helical structures, similar to what is observed for the entire protein(5, 15). The control peptide, however, does not show any tendency to form a helical structure, only a slightly more propensity to bend. Furthermore, it does not show any surface activity, contrary to the other peptides.

The specific intermolecular causes for the difference in the secondary structure formation and overall physicochemical behavior between the CAHS-inspired peptides and the SAHS-inspired control peptide remain elusive. However, there are similarities in the amino acid sequences of the CAHS-inspired peptides that are noteworthy: firstly, the CAHS-inspired peptides exhibit alternating sequences of positively charged amino acids (K and R) and negatively charged amino acids (E and D). This pattern resembles the polyampholytic sequences found in both natural and synthetic helical polypeptides (16). Secondly, the CAHS-inspired peptides contain a higher number of specific amino acids (M, A, L, E, and K) in their sequences compared to the control peptide. These particular amino acids are known to promote helical structures in peptide secondary structures(45). Among the model peptides, CAHS1M1 stands out as it has the highest occurrence of these amino acids: 10 out of 19 amino acids in CAHS1M1 are MALEK amino acids, occurring in two consecutive sequences of 6 (EAEAEK) and 4 (ELEK) residues each. The inherent structural attribute of CAHS1M1 could be a factor in its higher propensity to form a more helical secondary structure, even in the absence of desiccation-mimicking agents like TFE.

In summary, the model peptides chosen from the conserved region of CAHS protein upon drying undergo structural changes similar to the full-length proteins, known to be related to dehydration protection in tardigrades. Based on these results, we hypothesize that arginine is crucial to ensure the coil-to-helical transition of peptides inspired by CAHS. Since the conformational behavior under desiccating conditions is at the origin of the cryoprotective properties of CAHS, we suggest that model peptides designed to mimic tardigrade-proteins properties should contain this amino acid. Thus, this work offers promising sequences for the design of new synthetic peptide-based systems capable of mimicking the cytoprotective properties of dehydrated biologics and a generic rule for selecting specific sequences to use as a template. Our goal in the near future is to test whether novel synthetic model systems based on the CAHS model peptides presented here can mimic the cytoprotective properties of the naturally occurring tardigrade proteins.(4)

## ACKNOWLEDGMENTS

This work was supported by the Max Planck Graduate Center (MPGC) with the Johannes Gutenberg University Mainz. P.G.A. acknowledges the von Humboldt Foundation. I.M.I. acknowledges support from the Sectorplan Bèta & Techniek of the Dutch Government and the Dementia Research - Synapsis Foundation Switzerland. Y.R. acknowledges support from the China Scholarship Council (CSC).

