## Supplementary Material for "Structural adaptability and surface activity of tardigrade-inspired peptides"

### Supplementary Materials for:
### Structural adaptability and surface activity of tardigrade-inspired peptides

Giulia Giubertoni[1,||], Sarah Chagri[2,||], Pablo G. Argudo[2], Federico Caporaletti[1,2], Alessandro Greco[2], Leon Prädel[2], Alberto Pavan[1,5], Ioana M.Ilie[1,5], Yong Ren[2], David Ng[2], Mischa Bonn[2], Tanja Weil[2], and Sander Woutersen[1]

[1]Universiteit van Amsterdam, Amsterdam, The Netherlands
[2]Max-Planck-Institut für Polymerforschung, Mainz, Germany
[3]Vrije Universiteit, Amsterdam, The Netherlands
[4]Université libre de Bruxelles, Bruxelles, Belgium
[5]Amsterdam Center for Multiscale Modeling (ACMM), University of Amsterdam, P.O. Box 94157, 1090 GD Amsterdam, The Netherlands
[||]These authors equally contributed to this work.

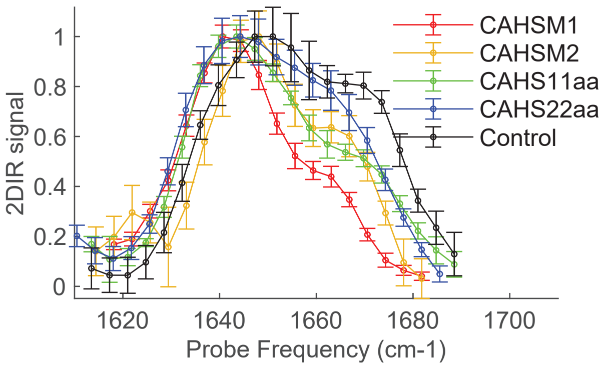


Fig. S1: Comparison between 2D-IR diagonal slices of the bleach signals of the isotropic 2D-IR spectra of the 5 different peptides.

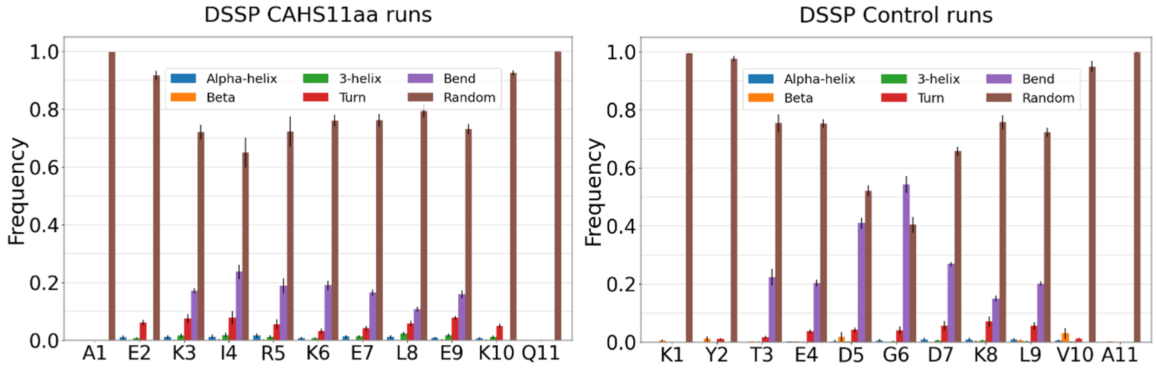


Fig. S2: Secondary structure probability per residue in CAHS11aa and Control peptide at physiological conditions.

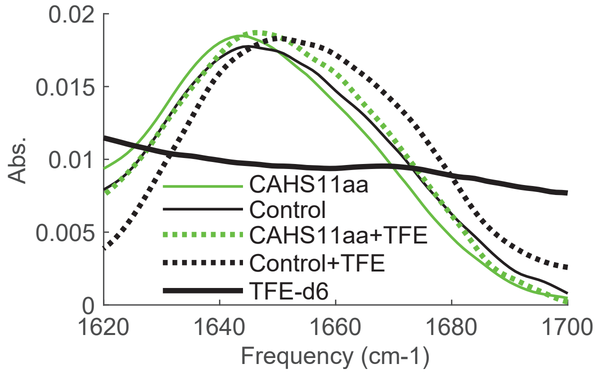


Fig. S3: Infrared spectra of CAHS11aa and control peptided at 0 and 50% TFE. We also report the IR spectrum of deuterated TFE (x 0.2), which also shows absorption bands in the amide I region. When subtracting the TFE IR spectrum to the peptide IR spectra, this overlap can cause problems leading to distortions in the background-corrected peptide spectra. These problems are overcome in 2D-IR since the TFE contribution to the 2D-IR spectrum is insignificant because 2D-IR signal scales as $\sigma^2$, and the $\sigma$ of the TFE absorption band is much lower than the ones of the amide I modes.

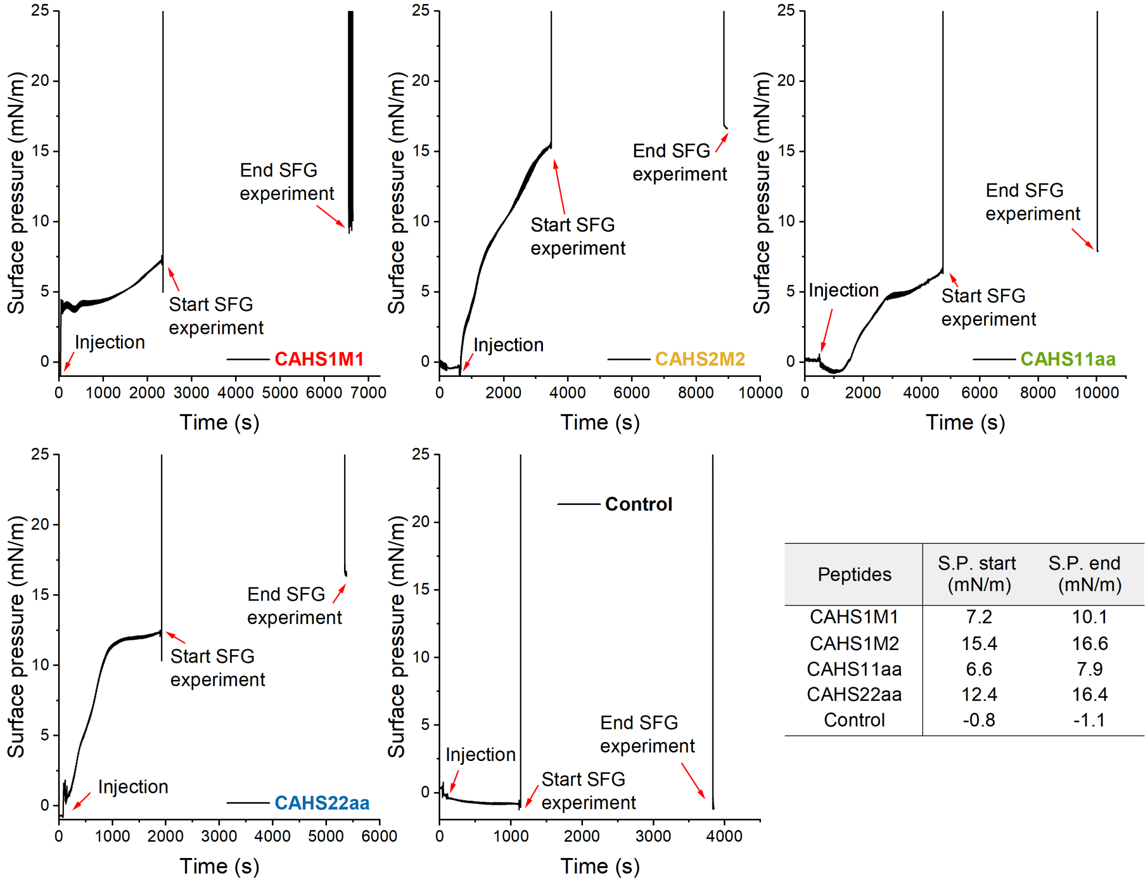


| Peptides | S.P. start (mN/m) | S.P. end (mN/m) |
|---|---|---|
| CAHS1M1 | 7.2 | 10.1 |
| CAHS1M2 | 15.4 | 16.6 |
| CAHS11aa | 6.6 | 7.9 |
| CAHS22aa | 12.4 | 16.4 |
| Control | -0.8 | -1.1 |

Fig. S4: Surface pressure of all peptides at 0.09 mg/ml in buffer solution and neutral pH. Note that the values were obtained before and after the SFG experiment was performed, which explains the missing values in this time range.

Table S1: Peak fitting parameters for SSP SFG spectra in Figure 4. $A_{NR}$ and $\varphi_{NR}$ are the amplitude and phase of the non-resonant signal, respectively. $A_n$ is the amplitude of the resonant signal, $\omega_n$ is the resonant frequency, and $\Gamma_n$ is the width of transition.

| | CAHS1M1 | CAHS1M2 | CAHS11aa | CAHS22aa | Control | Buffer |
|---|---|---|---|---|---|---|
| $A_{NR}$ | 0.047 | | | | | |
| $\varphi_{NR}$ | 1.385 | | | | | |
| A (HOH bend) | -8.8 | | | | | |
| $\omega$ (HOH bend) | 1658 | | | | | |
| $\Gamma$ (HOH bend) | 81.5 | | | | | |
| $\omega_1 (cm^{-1})$ | 1642 | | | | — | |
| $A_1$ | 7.1 | 1.4 | 0.5 | 7.2 | — | — |
| $\Gamma_1$ ($cm^{-1}$) | 39.2 | 32.6 | 22 | 26.7 | — | — |
| $\omega_2 (cm^{-1})$ | 1653 | | | | — | |
| $A_2$ | -0.3 | 7.6 | 6.9 | -0.9 | — | — |
| $\Gamma_2$ ($cm^{-1}$) | 11.5 | 52.4 | 42.9 | 12.1 | — | — |
| $\omega_3$ ($cm^{-1}$) | 1717 | | | | — | |
| $A_3$ | -3.9 | -3.5 | -4.4 | -4.4 | — | — |
| $\Gamma_3$ ($cm^{-1}$) | 31.4 | 27.4 | 32.5 | 38.1 | — | — |
